# Redefining the medicago sativa alphapartitivirus genome sequences

**DOI:** 10.1101/529610

**Authors:** Nicolás Bejerman, Debat Humberto, Verónica Trucco, Soledad de Breuil, Sergio Lenardon, Fabián Giolitti

## Abstract

In alfalfa samples analyzed by hightroughput sequencing, four *de novo* assembled contigs encoding gene products showing identities to alphapartitiviruses proteins were found based on BlastX analysis. The predicted amino acid (aa) sequences of two contigs presented 99-100% identity to the RNA-dependent RNA polymerase (RdRp) and the capsid protein (CP) of the recently reported medicago sativa alphapartitivirus 1 (MsAPV1). In addition, the remaining two contigs shared only 56% (CP) and 70% (RdRp) pairwise aa identity with the proteins of MsAPV1, suggesting that these samples presented also a novel *Alphapartitivirus* species. Further analyses based on complete genome segments termini and the presence/absence of alphapartitivirus RNA in several samples and public alfalfa RNA datasets corroborated the identification of two different alphapartitivirus members. Our results also likely indicate that the reported MsAPV1 genome was previously reconstructed with genome segments of two different alphapartitiviruses. Overall, we not only revisited the MsAPV1 genome sequence but also report a new tentative alphapartitivirus species, which we propose the name medicago sativa alphapartitivirus 2. In addition, the RT-PCR detection of both MsAPV1 and MsAPV2 in several alfalfa cultivars suggests a broad distribution of both viruses.

## Short communication

In Argentina, alfalfa (*Medicago sativa* L.) is a primary forage crop and a major feed component in dairy and beef cattle production systems. In 2010, we observed alfalfa plants showing symptoms of shortened internodes (bushy appearance), leaf puckering and varying-sized vein enations on abaxial leaf surfaces (Bejerman et al., 2011). Deep sequencing of alfalfa samples collected in central region of Argentina showing the dwarfism symptoms, revealed the presence of four RNA viruses: alfalfa mosaic virus (AMV), alfalfa dwarf virus (ADV), alfalfa enamovirus-1 (AEV-1) and bean leaf roll virus (BLRV) (Bejerman et al., 2015; 2016; Trucco et al., 2014, 2016) and one DNA virus: alfalfa leaf curl virus (ALCV) (Bejerman et al., 2018). Furthermore, four assembled sequences (contigs) analyzed by BlastX searches shared significant identity (E-value = 0), to the capsid and replication associated proteins encoded by alphapartitiviruses.

Partitiviruses particles are isometric and contain a double stranded RNA (dsRNA) bi-segmented genome; each segment contains a single open reading frame (ORF), which encodes a RNA dependent RNA polymerase (RdRP) or a coat protein (CP) (Nibert et al., 2014). Recently, the *Partitiviridae* family has been taxonomically reorganized; recently; it comprises five genera, *Alphapartitivirus*, *Betapartitivirus*, *Gammapartitivirus*, *Deltapartitivirus*, and *Cryspovirus* (Nibert et al., 2014). Partitiviruses corresponding to the same genus are usually reported in single infections; however they also have been described co-infecting a single host have also been described (Lesker et al., 2013; Lyu et al., 2018; Ong et al., 2017; Sabanadzovic and Valverde, 2011). In the latter case, it is a challenge to pair the two genomic segments to identify and annotate the genomes without mixing up genomic segments from different partitiviruses (Ong et al., 2017).

Alphapartitiviruses infect either plant or fungi. Plant-infecting alphapartitiviruses are associated with latent infections of their hosts and are transmitted with a high frequency via the ovule and by pollen to the seed embryo, but they cannot be transmitted by grafting or mechanical inoculation (Nibert et al., 2014). A new alphapartitivirus member infecting alfalfa, named as medicago sativa alphapartitivirus 1 (MsAPV1), was recently reported by analyzing public transcriptome dataset (Kim et al., 2018).

In this work we report two medicago sativa alphapartitiviruses, redefining the reported MsAPV1 genome sequence by Kim et al., (2018), and describing the medicago sativa alphapartitivirus 2 (MsAPV2) and redefined the reported MsAPV1 genome sequence. Furthermore, we detected both viruses in mixed infections in several alfalfa cultivars samples from Argentina and public alfalfa RNA datasets, suggesting a high prevalence of them.

In 2011, symptomatic alfalfa plants showing dwarfism were collected from a field located in Manfredi (Córdoba, Argentina), and total RNA was extracted using TRIzol Reagent (Life Technologies) according to manufacturer’s instructions. Total RNA was sent to Fasteris Life Sciences SA (Plan-les-Ouates, Switzerland), where the bands located between 21 and 30 bp were excised, purified, processed and sequenced on an Illumina HiSeq 2000. The obtained raw reads were filtered by quality and trimmed using the FASTX-Toolkit as implemented in http://hannonlab.cshl.edu/fastx_toolkit/index.html. The remaining short reads were de novo assembled using Velvet 1.2.10 with the velvet optimiser parameter available at https://www.ebi.ac.uk/~zerbino/velvet/. The obtained contigs were subjected to bulk BlastX searches (E-value < 1e-5) against a local database of the non-redundant virus proteins. Four contigs ranging from 1,650-1,840 nt and sharing high similarity (56-100%) with alphapartitivirus CP and RdRp genes were retrieved, further inspected and annotated as described in Debat and Bejerman (2018). The contigs were curated by mapping the filtered sequencing reads using the Geneious 8.1.9 (Biomatters, inc) mapper with low sensitivity parameters and assessing the consensus by hand. The curated contigs were initially designated as alphaA1 (1,872 nt, mean coverage 31.6X, supported by 2,782 reads, encoding an RdRP); alphaA2 (1,687 nt, mean coverage 11.5X, supported by 876 reads, encoding a CP); alphaB1 (1,845 nt, mean coverage 38.9X, supported by 3,357 reads, encoding an RdRP); and alphaB2 (1,714 nt, mean coverage 28.5X, supported by 2,314 reads, encoding a CP). AlphaA1 and alphaA2 shared 98.7% and 99.8% nt identity with the reported MsAPV1 dsRNA1 (MF443256) and dsRNA2 (MF443257), respectively. whereas alphaB1 and alpha B2 shared 68.9% and 55.6% nt identity with the reported MsAPV1 dsRNA1 (MF443256) and dsRNA2 (MF443257), respectively. Thus, we tentatively grouped the identified contigs into two putative viruses corresponding to MsAPV1 (alphaA1/alphaA2) and MsAPV2 (alphaB1/alphaB2). However, evidence presented below based on the complete genome segment sequences and RNA levels indicated that the genome segments could be paired differently. So, alphaA2 (almost identical to the reported MsAPV1 dsRNA2 (MF443257)) is in fact the RNA2 of MsAPV2, and thus alphaB2 should be paired with alphaA1.

To obtain the complete sequences of these putative alphapartitivirus genome segments, their terminal sequences were amplified by 5’and 3’rapid amplification of cDNA ends (RACE) that was carried out as described previously by Phelan and James (2016) which resulted in assembly of full genome sequences that were deposited in the GenBank database under accession numbers MK292286 (MsAPV1 dsRNA1), MK292287 (MsAPV1 dsRNA2), MK292288 (MsAPV2 dsRNA1) and MK292286 (MsAPV2 dsRNA2). Primers used to amplify the terminal regions are listed in Supplementary Table 1.

**Table 1.**
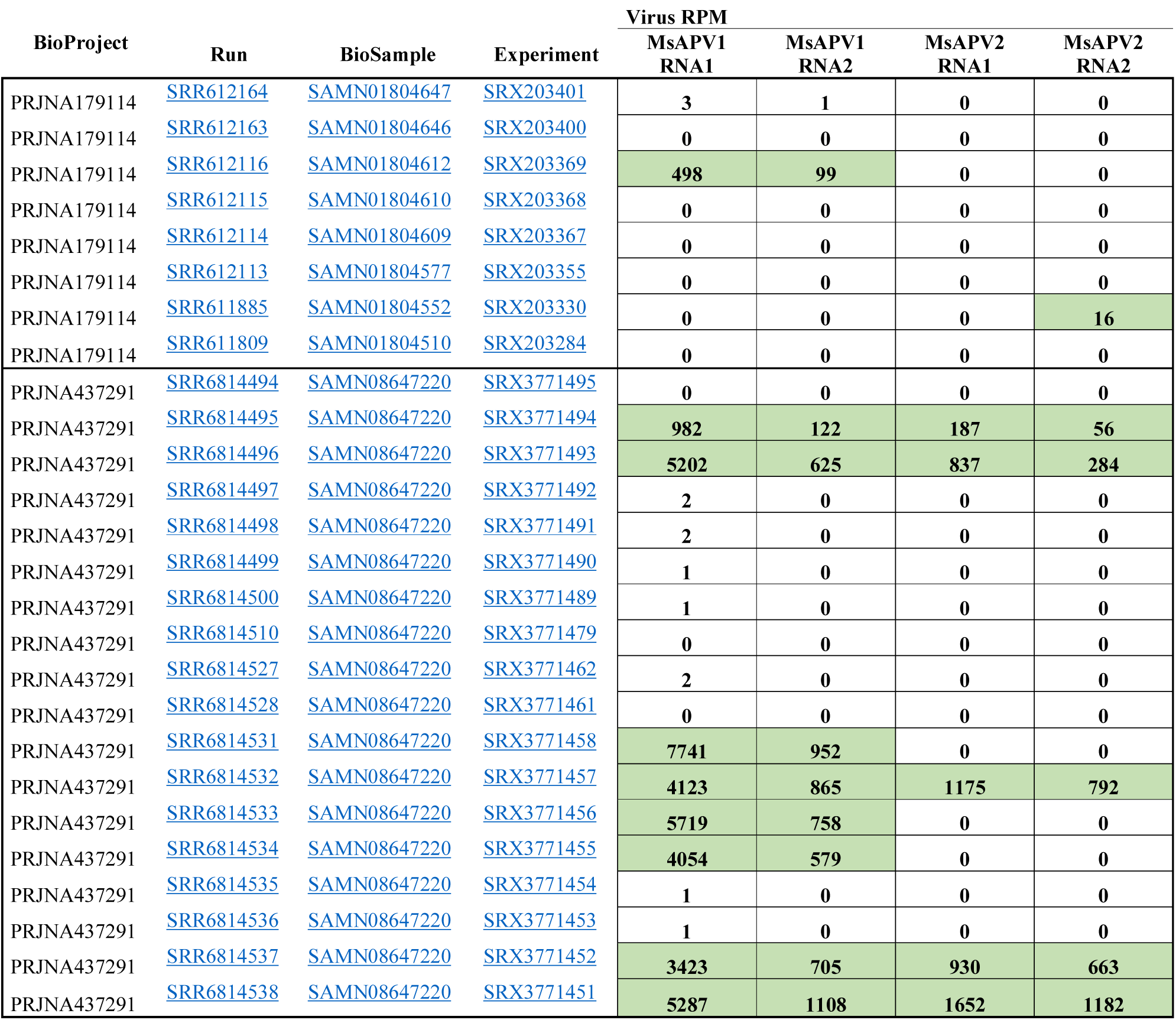
Virus RNA levels corresponding to each genome segment of MsAPV1 & 2 in multiple public datasets. Values express virus derived reads per million of total reads (RPM) based on mapping to the assembled genome sequences. An arbitrary (and conservative) cut-off of 5 RPM was considered to indicate virus RNA presence, which is indicated in green cell coloring.

The complete sequences of the revisited MsAPV1 dsRNA1 (extended from the formerly designated alphaA1 contig) and dsRNA2 (formerly designated alphaB2 contig) were determined to be 1942-and 1806-nt long, respectively (Fig. 1A) with GC contents of 38.62% and 43.79%, respectively; whereas the complete sequences of the novel MsAPV2 dsRNA1 (alphaB1) and dsRNA2 (alphaA2) were determined to be 1939- and 1764-nt long, respectively (Fig. 1A) with GC contents of 43.73% and 44.72%, respectively. GC content is likely similar to those values reported for other partitiviruses (Phelan and James, 2016). Each dsRNA contains a single open reading frame (ORF) with 5’ and 3’untranslated regions (UTRs). Interestingly, multiple alignments of 5’UTRs revealed that these sequences are highly conserved, being the first 9 nt at the 5’terminus of the dsRNA1 and dsRNA2 fragments of MsAPV1 identical, whereas the first 10nt at the 5’terminus of the dsRNA1 and dsRNA2 fragments of MsAPV2 also are identical, but different from the 5’ termini of the MsAPV1 segments (Fig. 1B). Furthermore, the 5’UTR of MsAPV1 and MsAPV2 dsRNA1 and dsRNA2 were predicted to fold into stem-loop structures, when examined using Mfold software (http://mfold.rna.albany.edu) (Fig. 1C). Similar stem-loop structures, which are likely to play an important role in dsRNA replication and virion assembly, have been described in other partitiviruses (Guo et al., 2017; Lesker et al., 2013; Osaki and Sasaki, 2018).

**Fig. 1.**
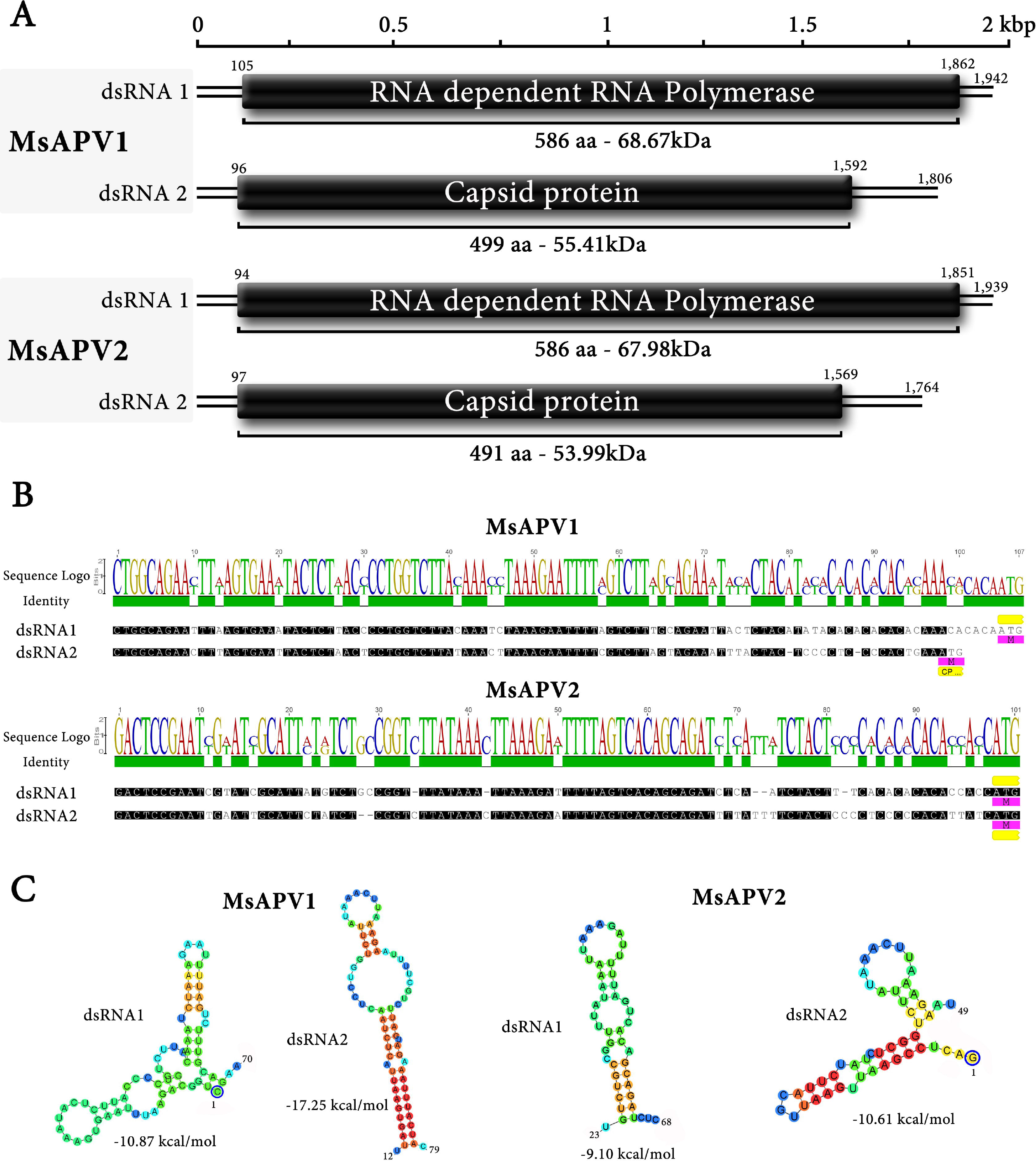
**A**. Genomic organization of medicago sativa alphapartitivirus 1 and 2. The numbers above the solid lines indicate the nucleotide positions of the beginning and end of the 5’- and 3’-UTRs. **B**. Multiple alignments between 5’UTR of dsRNA1 and 2 of MsAPV1 and MsAPV2. **C**. Potential secondary structures formed by 5′-terminal sequences of positive strands of two segments of MsAPV1 and MsAPV2.

Sequence analysis of MsAPV1 and MsAPV2 complete dsRNA1 revealed that it contained a single ORF encoding a putative 586-amino acid (aa) RdRP protein, where a RdRp conserved domain was found (cd01699, e-values =3.73e−05 and 3.40e−07). Moreover, the multiple-protein alignment of the deduced aa sequences of the RdRps of MsAPV1, MsAPV2 and plant-infecting alphapartitiviruses confirmed that this protein includes the six conserved motifs (III to VIII) described for partitiviruses (Liu et al., 2015). Sequence analysis of MsAPV1 and 2 dsRNA2 revealed that it contained a single ORF encoding a putative 499- and 491-aa CP protein, respectively.

Pairwise comparison using Sequence Demarcation Tool (SDT v1.2) (Muhire et al., 2014) showed that at aa level the RdRp of the redefined MsAPV1 is 100% identical to that one encoded by the reported MsAPV1 (Fig. 2A), whereas its CP is just 56% identical to that one of the redefined MsAPV1 reported MsAPV1 (Fig. 2B). On the other hand, the RdRp of the MsAPV2 is just 70% identical to that one encoded by the redefined MsAPV1 and the reported MsAPV1 (Fig. 2A), whereas its CP is 99% identical to that one of the alphapartitivirus 1 and the former MsAPV1 (Fig. 2B), which likely suggest that MsAPV1 genome segments of the reported MsAPV1 could be paired differently that previously described. Both the redefined MsAPV1, and MsAPV2 have a degree of identity below 90% and 80% for RdRp and CP protien, respectively, when compared with other partitiviruses, respectively, which is the species demarcation threshold recently recommended recently for alphapartitiviruses (Vainio et al., 2018).

**Fig.2.**
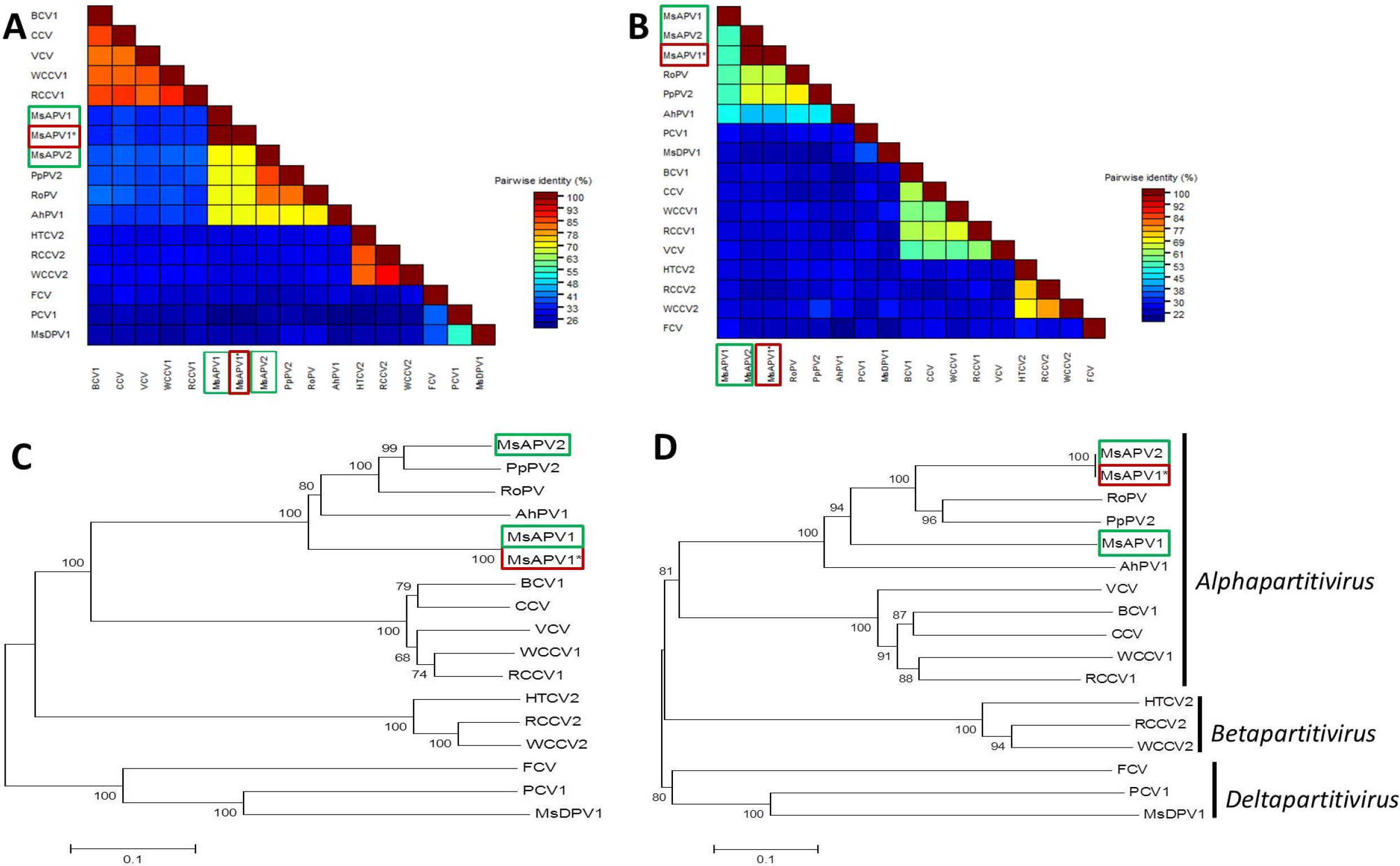
**A and B**. Amino acid percentage pairwise identities between MsAPV1 and MsAPV2 RdRp (**A**) and CP (**B**) with those from selected partitiviruses members calculated using SDT v1.2 (Muhire et al., 2014). **C.** Maximum likelihood phylogenetic tree of MsAPV1 and 2 and selected partitiviruses members based on the RdRp amino acid sequences. **D**. Maximum likelihood phylogenetic tree of MsAPV1 and 2 and selected partitiviruses members based on the CP amino acid sequences. All trees were constructed using MegaX software. LG + G model was used to construct the phylogenetic tree based on RdRp amino acid sequences; whereas model LG + G + F was used to construct the phylogenetic tree based on CP amino acid sequences. Bootstrap values out of 1000 replicates are given at the nodes. The bar below each tree represents substitutions per site. Accession numbers of every virus used to perform SDT analysis and construct phylogenetic trees are listed in Supplementary Table 2.

To further assess our speculation that the reported MsAPV1 segments derived from two different viruses we analized to the raw data which was the source of Kim et al (2018) study, which pooled eight libraries before de novo assembly. Raw reads corresponding to RNAseq NGS libraries (SRA: SRP017117), associated to NCBI Bioproject PRJNA179114 and additionally from an independent study oriented to elucidate the molecular genetic basis of herbivory between butterflies and their host plants, including alfalfa (Nallu et al 2018, PRJNA437291, SRA: SRP134094), were analyzed according to Debat and Bejerman (2018). Briefly, the respective public datasets were downloaded and reads of each library including our NGS data were mapped against the assembled virus genomes sequences using Bowtie2 v2.3.4.3 available at http://bowtie-bio.sourceforge.net/bowtie2/ with default settings and the fast end-to-end preset. The obtained values were normalized as mapped reads per million (RPM) for each library. Interestingly, we observed a differential pattern of presence and absence of the diverse alphapartitivirus segments in each library (Table 1). When we analyzed in detail the transcriptome data used by Kim et al., (2018) we observed that two of the eight pooled libraries presented virus RNA derived from both segments of the revisited MsAPV1 alphaA1 and alpha B2, and only one library harbored virus reads of segment 2 (alphaA2) of MsAPV2 (which was previously annotated as segment 2 of MsAPV1). Thus, we believe that the use of a pooled transcriptome dataset for virus discovery could misrepresent the virus landscape associated to each specific sample that led to a miss assignment of alphapartitiviruses genome segments. Moreover, when we explored 18 alfalfa RNAseq datasets from bioproject PRJNA437291 we observed that in three libraries only virus RNA from both genome segments of the redefined MsAPV1 (alphaA1 and alpha B2) were found, while in five additional runs we detected virus RNA from both segments of MsAPV1 (alphaA1 and alpha B2) and MsAPV2 (alphaA2 and alpha B1), suggesting mixed infections of alphapartitiviruses in those samples (Table 1). The detection of only both segments (alphaA1 and alpha B2) of the redefined MsAPV1 in several samples likely indicates that the former MsAPV1 genome could be revisited.

Following sequence annotation and the assessment of RNA levels, the redefined MsAPV1 and 2 predicted RdRp and CP aa sequences were aligned with the corresponding sequences of selected partitiviruses, described in Supplementary Table 2, using MUSCLE (Edgar, 2004) as implemented in Mega X (Kumar et al., 2018). ML phylogenetic trees were inferred using Mega X (Kumar et al., 2018) with LG + G and LG + G + F aa substitution models, respectively, and 1000 bootstrap replicates. Phylogenetic analyses supported the pairwise comparison results showing that the redefined MsAPV1 RdRp grouped within alphapartitiviruses with the reported MsAPV1 whereas MsAPV2 RdRp clustered with those encoded by pyrus pyrifolia partitivirus 2 (PpPV2) and rose partitivirus (RoPV) (Fig. 2C). On the other hand, the revisited MsAPV1 CP formed a monophyletic cluster within alphapartitiviruses whereas MsAPV2 CP grouped with those encoded by the reported MsAPV1, PpPV2 and RoPV (Fig. 2D). This suggest that the redefined MsAPV1 and MsAPV2 do not share a common immediate ancestor, so it is tempting to speculate that they originated from two distinct ancestral partitiviruses and adapted to the same host.

Recently, it was described the presence of the reported MsAPV1 in alfalfas from USA (Nemchinov et al., 2018). Given that the presence of alphapartitivirus RNA sequences in that study was assessed by RNAseq reads mapping using as reference the published sequences by Kim et al., (2018), it is tempting to suggest, based on our results, that these alfalfas could be infected with both MsAPV1 and MsAPV2.

It was reported that there is higher sequence identity between the corresponding 5′ UTRs of CP and RdRp segments belonging to the same alphapartitivirus than those belonging to different species (Ong et al., 2017). Therefore we verified if genomes were consistently annotated in this study by comparing the 5′ UTRs of segments of the redefined MsAPV1 and MsAPV2. The comparison revealed a 70.5% and 70.1% identity between the 5′ UTRs of CP and replicase segments of the redefined MsAPV1 and MsAPV2, respectively. Whereas identities values when 5′ UTRs of CP and replicase segments of the redefined MsAPV1 were compared against those of MsAPV2 were lower than 60%. This supports our tentative pairing of the revisited MsAPV1 and MsAPV2 genomes in this study. Plant-infecting partitiviruses are efficiently pollen/seed transmitted, but are cryptic, so infected-plants are symptomless (Nibert et al., 2014), therefore they are not considered as quarantine pest and they are not tested when alfalfa seeds are traded. Consequently, both the redefined MsAPV1 and MsAPV2 could have been transmitted to many alfalfa cultivars inadvertently, resulting in their dissemination all around the world through the exchange of alfalfa seeds, which led to its detection in diverse contexts. In this regard, in order to assess a tentative prevalence landscape of alphapartitivirus in alfalfa, we studied the occurrence of the revisited MsAPV1 and MsAPV2 by RT-PCR assays in leaf tissue of several alfalfa cultivars, using two set of virus-specific primers designed from dsRNA1 (Supplementary Table 1) in order to detect the redefined MsAPV1 and MsAPV2 in alfalfa leafs. Interestingly, we detected both dsRNA1 sequences in all samples tested, suggesting that the presence of both viruses as mixed infections is common in alfalfa (Supp. Fig. 1).

In summary, we describe the complete genome sequences of two alphapartitivirus infecting alfalfa, revisiting the reported MsAPV1 genome sequence by Kim et al., (2018) and naming the second one as medicago sativa alphapartitivirus 2. Both the revisited MsAPV1 and MsAPV2 were found in mixed and single infections in alfalfa samples from diverse regions of the world and should be considered as new members of the genus *Alphapartitivirus*. We also implemented here a simple and valuable tool based on RT-PCR detection to screen for the presence of both viruses in alfalfa cultivars. Our study provides new avenues to explore and confirm virus genome segments when partitiviruses (or any multi-segmented viruses) are co-infect a sample, which appears to be genuinely common.

## Acknowledgments

We thank Hector Tharghetta (Barenbrug Palaversich) for providing seeds of the alfalfa cultivars evaluated in this study and Monica Cornacchione (EEA-INTA-Santiago del Estero) for growing the alfalfa cultivars evaluated in this study.

## Compliance with ethical standards

### Ethical Approval

This article does not contain any studies with human participants or animals performed by any of the authors.

### Funding

This work was partially supported by the PNPV 1135022 project of INTA.

## Supporting information

MsAPV1 and MsAPV2 virus sequences

