## Supplementary Table 1 for "Redefining the medicago sativa alphapartitivirus genome sequences"

Supplementary Table 1. List of primers used to validate infection and obtain the complete genomes of MsAPV1 and MsAPV2

| Primer Name | Sequence | PCR product (bp) | Purpose |
| --- | --- | --- | --- |
| Ms1A3R | CTATGCCATATCGTGAAGAAC | 451 | MsAPV1 RNA1 3´RACE |
| Ms2A3R | GACGAACTACAACTATTAGC | 445 | MsAPV2 RNA1 3´RACE |
| Ms1B3R | CACTGTTCAATGTTACTGC | 507 | MsAPV1 RNA 2 3´RACE |
| Ms2B3R | CTGCCAATCAGATTACTGC | 521 | MsAPV2 RNA2 3´RACE |
| Ms1A5R | TCCGAGTGTGTTCGATAGC | 378 | MsAPV1 RNA1 5´RACE |
| Ms2A5R | GTACAGGCCTAAGTTTCTC | 403 | MsAPV2 RNA1 5´RACE |
| Ms1B5R | GCTCCATTTGCTGGAATAC | 399 | MsAPV1 RNA2 5´RACE |
| Ms2B5R | GTAGATACAGTCTTCCTAGC | 410 | MsAPV2 RNA2 5´RACE |
| 1FD | CTCAAGTGGTCATGATCTC | 674 | MsAPV1 detection |
| 1RD | GAGTAATGAAAGGATAGCAC |  |  |
| 2FD | AAGACTAACGACCATGGAC | 848 | MsAPV2 detection |
| 2RD | GCTAATAGTTGTAGTTCGTC |  |  |
