## Supplementary Table 2 for "Redefining the medicago sativa alphapartitivirus genome sequences"

Supplementary Table 2. Accession number of each virus used to perform pairwise comparisons and construct the phylogenetic tree

| Acronym | Full Name | Accession number |
| --- | --- | --- |
| MsAPV1* | medicago sativa alphapartitivirus 1 | MF443256/7 |
| PpPV2 | pyrus pyrifolia partitivirus 2 | LC221826/7 |
| RoPV | rose partitivirus | KU896858/9 |
| VCV | vicia cryptic virus | AY751737/8 |
| AhPV1 | arabidopsis halleri partitivirus 1 | LC151461/2 |
| WCCV1 | white clover cryptic virus-1 | AY705784/5 |
| RCCV1 | red clover cryptic virus 1 | KF484724/5 |
| BCV1 | beet cryptic virus 1 | EU489061/2 |
| CCV | carrot cryptic virus | FJ550604/5 |
| HTCV2 | hop trefoil cryptic virus 2 | JX971980/1 |
| WCCV2 | white clover cryptic virus 2 | JX971976/7 |
| RCCV2 | red clover cryptic virus 2 | JX971978/9 |
| MsDPV1 | medicago sativa deltapartitivirus 1 | MF443258/9 |
| PCV1 | pepper cryptic virus 1 | JN117276/7 |
| FCV | fig cryptic virus | FR687854/5 |
