## Supplementary Figure 1 for "Redefining the medicago sativa alphapartitivirus genome sequences"

Supplementary Figure 1. Agarose gel electrophoresis showing the results of the RT-PCR detection of MsAPV1 and MsAPV2 in several alfalfa cultivars. The order of alfalfa samples is: cvs “Monarca” (lane 2), Antares (lane 3), SARDI 10 (lane 4), Sardi 7 (lane 5), Sardi Grazer (lane 6), Pegasis (lane 7), Soraya (lane8), Genesis II (lane 9), Traful (lane 10). Water control (W) is in lane 1 and 100 pb DNA ladder (Biodynamics) is in lane M.

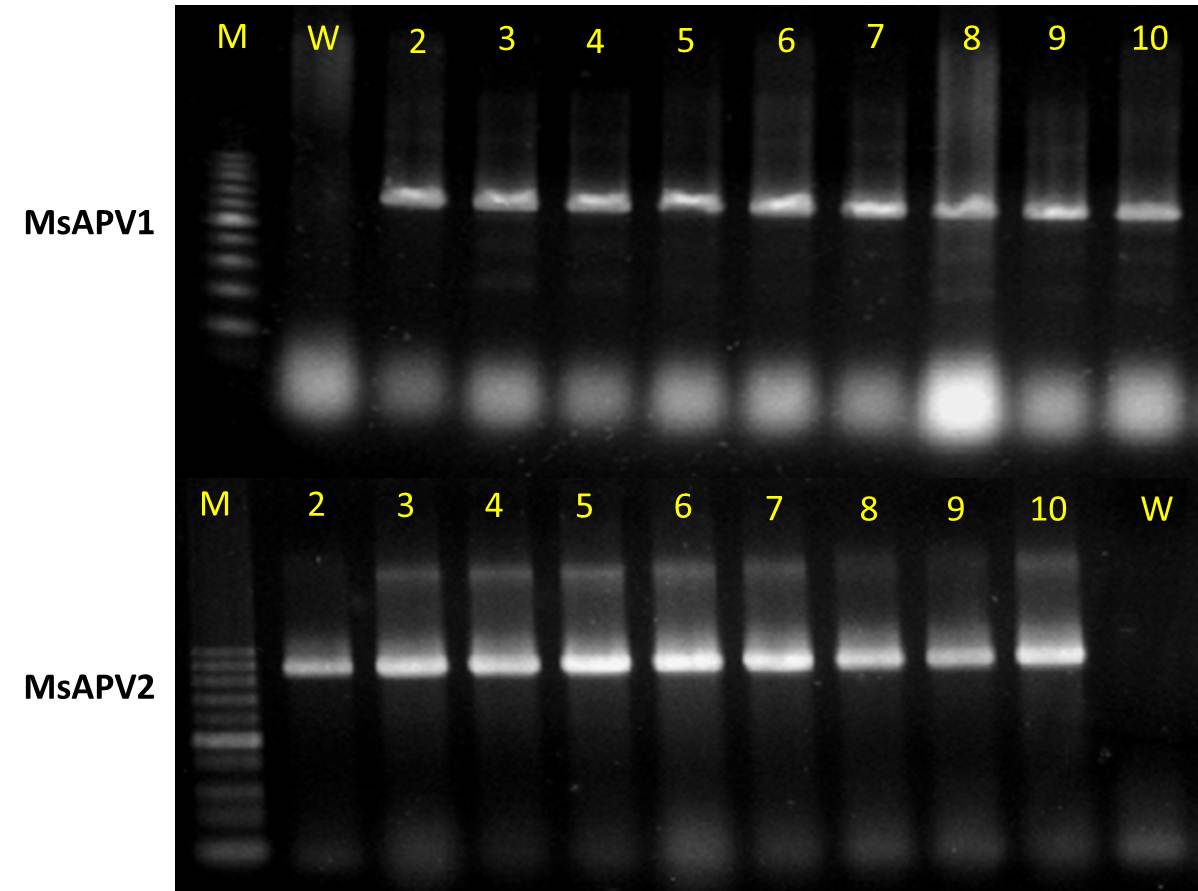
