## Supplementary material for "Redefining the medicago sativa alphapartitivirus genome sequences": MsAPV1 and MsAPV2 virus sequences

>MsAPV1 RdRp dsRNA1 1942nt

CTGGCAGAATTTAAGTGAAATACTCTTACCCCTGGTCTTACAAATCTAAAGAATTTTAGTCTTTGCAGAATTACTCTACATATACACACACACACAAACACACAATGAAGAATATCGAGATAATCGGTTACAAGCCCTCTTTGGCTAAACCTATTCGTGGTAATGTTGATCCTAACTCAAACATTAATTACGGTAATATCGTCGATTATGCTTTAAGAAAATACCTCACTAATGAGGAGTTTACTATCGTCACACGTGGCTATCGTCGCTCACAATGGGCTGAAGACAGTCTTAAAAGTGATTTAGATAAACTAGATTCAGACTACTTTCCCGTGCTCAAAGATACCCATTATTATAATGCTATCGAACACACTCGGAAATTATTTAAACCTGATTCTTTACTTAAGCCTATACATTTTTCTGACTTGCGCCATTACCCATGGCAGTTATCTACTAATATCGGTGCCCCTTTTGCTACTAGTAAAAGTTGGAACGAATACGTAATACAGAAATTCGATGAAGGTTTTACCAAATCTTATTATCGAGACTTATTTCGTGAAGCCCACGGAGAAAGTCTACTTCCTGAGATGATTGATCGTCGTATGACGAAACGCAATCTATACAATGAAATGTTCTTTATTAACCGCACAAATATCCACTTAATTAAAGATGGACACACTACTAACTCAAGTGGTCATGATCTCAAATATTGGAACACAGCATTTGCTCGACAGCATCTAGTTGAATCCCATGATGAAGATAAGATCCGCTTAGTATTCGGTGCTCCTTCCACATTTTTGATGGCTGAACTGACGTTTATCTGGCCTTTACAGACAAGCTTACTATACCAAGGAGAACGTTCTCCTATGCTATGGGGCTATGAAACCACTACTGGTGGTTGGTCTAGACTATATAAATGGGCCTCCTCTAGGATGCCTAGGTATGATTTTGTTGCTACTCTCGACTGGAAACGGTTTGATAGAGACGCTAGACATACTGTTATCTCTGATATACATCAACTTGTTATGAGATCATATTTTGATTTCAATAATGGATATCATCCTACAATCCACTACCCTGACTCCACCGGAGCCAACCCTCAAAGAATTGAGAACCTATGGAATTGGATGACTGACGCAACTTTAACGATTCCGCTAATGTTGCCTGACGGTAAGATACTTCGATTTAAACATTCTGGTATATACTCTGGATACTTCCAGACACAAATACTAGATTCGATGTACAATTGTGTTATGATCTTCACTGTTCTATCTAGAATGGGGTTTGATCTTGAAAGAGTTGAAATTAAAGTACAAGGAGATGATTCCATTTTCTTAATGTGCTATCCTTTCATTACTCTTCAAAATACGTTTCTACAGATGTTCGCTCACTATGCGAAAATTTACTTTGGATCAACTTTAAATATTGATAAAAGTGAAATATTACCTAGCCTTGAAAACGCTGAGGTATTGAAATATCGCAACCATGGTACTATGCCATATCGTGAAGAACTACAACTATTAGCTATGTTGAGACACCCTGAAAGGACTGTCTCGTTACCATCGCTGATGGCTCGATCCATCGGCATTGCTTATGCTAACTGTGGATTCCACTCTCGTGTCTACCAAATTTGCGAGGATATCTATAACTTTCTGAAAGCCGGAGGCTATAGCCCTGACCCGCATGGTTTACCAGGTAGTCTACGCTATAGGCAAAATTACGTTCCAGGATATTCTGAAGTTGACATTAGTCACTTCCCTAGTTATTTTGAAACTGTCCGCCTTCTCCAAGAGCCAACTCGTGACTTAGTTTCCGAAAAGCATTGGCCATTAAAACACTTTATCGGTATCCCCGGAAAGTCTTAAGTTTAAGACGTTATTTAGGCGTCACCTTAAGTTAATAAAATAAAAGGAAACCCCAATCCGAAAAAAAAACAAAAGAT

>RdRp 586aa

MKNIEIIGYKPSLAKPIRGNVDPNSNINYGNIVDYALRKYLTNEEFTIVTRGYRRSQWAEDSLKSDLDKLDSDYFPVLKDTHYYNAIEHTRKLFKPDSLLKPIHFSDLRHYPWQLSTNIGAPFATSKSWNEYVIQKFDEGFTKSYYRDLFREAHGESLLPEMIDRRMTKRNLYNEMFFINRTNIHLIKDGHTTNSSGHDLKYWNTAFARQHLVESHDEDKIRLVFGAPSTFLMAELTFIWPLQTSLLYQGERSPMLWGYETTTGGWSRLYKWASSRMPRYDFVATLDWKRFDRDARHTVISDIHQLVMRSYFDFNNGYHPTIHYPDSTGANPQRIENLWNWMTDATLTIPLMLPDGKILRFKHSGIYSGYFQTQILDSMYNCVMIFTVLSRMGFDLERVEIKVQGDDSIFLMCYPFITLQNTFLQMFAHYAKIYFGSTLNIDKSEILPSLENAEVLKYRNHGTMPYREELQLLAMLRHPERTVSLPSLMARSIGIAYANCGFHSRVYQICEDIYNFLKAGGYSPDPHGLPGSLRYRQNYVPGYSEVDISHFPSYFETVRLLQEPTRDLVSEKHWPLKHFIGIPGKS

>MsAPV1 CP dsRNA2 1806nt

CTGGCAGAACTTTAGTGAATTACTCTAACTCCTGGTCTTATAAACTTAAAGAATTTTCGTCTTAGTAGAAATTTACTACTCCCCTCCCCACTGAAATGGCTTCTAATTCAGGTACCAAATCTGATGAGGTTCAAAAGAGCCTCCAGACTCTGAAATCCTTAATAGATGAGAACAAGATTGATCAACAGCTCCTACAAACTCTAGGCCTGAAATCACTCGACAAATTCTCTCCTACTGAGGTTGAAGACCCATTTCCCTCTAAAGCTACGGTTGTAGCTCAGACAACACTATCTGCCGATAGGCGCACGCCTAAAGAGCAAGATAAGTCACTTGCTGATGCTGTTGAGCCAAAGGCTCCTACTGTCGCTTCTGATATTCTAGCTCCATTTGCTGGAATACATATTACTTATGCTCCTAGATTGAGCACGAGTAGTTACTCCCCCTCCTCTTTGATGATGGACTATATCGTTCATCAAATCAACTCTAACTTAGTTGATAATTTTTATTTCAAAAGGACCTCCCCTGATTATCATCCCTACATCATCCGCCTATATTATGGCGTGATCTTCTGGGTTCAGTGCATTCGCGCTGGACACTATGTTGGTGAACTTGATATGAGTAAGCACCAATTTCTGGTCCGCTTCCTAGATGCTCATCCACTTGAAAGCCTGACCGTCGCTGGTCCCCTCATCCCTCTGTTCAAAACGCTGTGTGCTTCACAGCCTGAAATACCCACTTTCGGGAAAGTCTATCCAAGGCTACCCGCAGTTGTCGGCCCTAACCGCCGTGATGAATTTATTAAGAATGATATCACTTCAAACCTGCTTCCAAACGTCCCTGGGATTTTTGCGTTGCTTGAGCACCTTAATGGTATCATCAACCCCACCCCTCCCGCTGACCCCGCTTACCCAAAGAAGGGTATGCACATACCTGTTACTGCTACTACCAACCAGGCGACCATTTTTGGTCATCACACTTTTCCTGTGCCTGCTGAGAGAACTAACAGAGACCGTTGGGCTCTTTGCTCTTCTGGTTTACAATATGAATGTGAAGCTGATGCCCGTCTGAATGAGGCCTTCGCTGAAAGGTACTCAAAATTCAGATTTCCTACTCATAGGGCAAATGATGCACTTATTGAGATAGATTACTTTCTATCTATGGATGTATCAATGGCTTGGTTCGCCCAAGTCAAGAAAGTTGCTGCTGCTGCCGCTGCCTATTTTGAAGGGTCTGGCACCCTCGTTGACTGTCCTCCTCATGGAATAGCCGCTAACCAGATTATCGTTAGTTATGTTCCACCACTGTTCAATGTTACTGCCCCTACCCGATCTGCAGACCCTGCCTCCCAGTTCCCGTTTGCTTTCAAACTGGCAACCTCCGCCCGCAACCTTCCCACTCTTTCCGAATCAATGGCCGCTATGGCCCAAACTAATGTGATGATGTATCGCACACATCCTTATTTTGGTGACTTTGGAATTGAGACTATCGAAGGTGAATTTTGGGATATTCGCCCCCTTGAATCTTCTTTAACTGATTCTTCCACTTACCTGTCACTTGATGACTCGGTAACTAAGATGATGAAGTCTAAGAATTAGATTCTGCGTTTCCGTCGTCTAGATGTTTCTAGTTCCGGTTTTATTTCGCTTCTTTATTTTGTTAGGATTTAGAACAAAAACCAAAAAAAATTTTAAAATCACTTGCTGTTTTTTTATTTCTCGTCCTGCGTTTTTCCTTTTGTCAGTCAATTAAGGCTGACCGATACGATCATCACACCCAAAATTGGTCAAAAAAAAAAAAAAAAATATC

>CP 499 aa

MASNSGTKSDEVQKSLQTLKSLIDENKIDQQLLQTLGLKSLDKFSPTEVEDPFPSKATVVAQTTLSADRRTPKEQDKSLADAVEPKAPTVASDILAPFAGIHITYAPRLSTSSYSPSSLMMDYIVHQINSNLVDNFYFKRTSPDYHPYIIRLYYGVIFWVQCIRAGHYVGELDMSKHQFLVRFLDAHPLESLTVAGPLIPLFKTLCASQPEIPTFGKVYPRLPAVVGPNRRDEFIKNDITSNLLPNVPGIFALLEHLNGIINPTPPADPAYPKKGMHIPVTATTNQATIFGHHTFPVPAERTNRDRWALCSSGLQYECEADARLNEAFAERYSKFRFPTHRANDALIEIDYFLSMDVSMAWFAQVKKVAAAAAAYFEGSGTLVDCPPHGIAANQIIVSYVPPLFNVTAPTRSADPASQFPFAFKLATSARNLPTLSESMAAMAQTNVMMYRTHPYFGDFGIETIEGEFWDIRPLESSLTDSSTYLSLDDSVTKMMKSKN

>MsAPV2 RdRp dsRNA1 1939nt

GACTCCGAATCGTATCGCATTATGTCTGCCGGTTTATAAATTAAAGATTTTTAGTCACAGCAGATCTCAATCTACTTTCACACACACACCACCATGCGAAACACAACCGTTATTGGGCACAAGCCCAGTCTTGCTAGGCCTCTTTATGGAAATCCAGATGCTGGATCCAATCCTGCCTACGCTGATACAGTTGATCACGCACTACGTCGATATTTAACTCCTGAAGAATTCAACAGAGTTGTTAATGGCTATCGCCGCTCCCCATGGAACGAAGATGCACTAAATGATGATATCGCTAAATTAAACAGCGATGAACATACCGTTATCAAGGATGAGCACTATTATAAAGCCATCGAACACACCAAGAAACTCTTCACACCAAAGGAGAAACTTAGGCCTGTACACTTTACAGACCTTCGCCACTATCCATGGCAACTTGCCTCCAACATAGGCGCCCCTTTTGCCTCCAGCAAAGAATGGCAGCATTATGTCAATGACAAGTTCCTCAAAGGCCATACCGCTACTGAGGTTCGCGACCTATTTAAGGAAGCACATGGACAATCTCTTGAACCAGAGATCATCGATAGACGTATGACTAAGCGTAATCTATACAACGAGATGTTTCTCATTAACCGAACGAACATCCATCTTATAAAAGATGGACGTAAGACTAACGACCATGGACATGATTTGCGATACTGGAATACAGCATTCGCCCGTCAACACCTGGTTGAAGATGATGAACCTGATAAGGTCCGTCTAGTCTTCGGCGCCCCGTCAACTTTGCTAATGGCAGAGTTAATGTTTATATGGCCCATACAAGTCAGCTTGCTGAACCGTGGACCTGAATCTCCAATGCTTTGGGGATATGAAACTATTACTGGCGGATGGTCCCGTCTTCACACATGGGCTTCTACAGCCCTTCCTCGATTTGAGTCAGTAGTGACTCTAGATTGGAGCCGATTTGATAAAGATGCACGCCACACTGTCATTCGTGATATTCACGCAATGATTATGCGACCGATGTTTACTTTTGAACATGGCTACCACCCTACCTACCACTACCCTGACACTTCCGACACCAACCCTGCGAGGTTAGAGAACTTATGGAATTGGATGACTGATGCCATCCTCACCACCCCTCTAATTCTACCAGATGGCAGTGTCCTAAGATTTAACCATTCAGGTATTTACTCTGGATATTTCCAGACTCAAATATTAGACTCTATTTACAATTGTGTAATGATTTTTACGATACTATCCCGTATGGGATTCGACTTAGATCGCGTAGAAATTAAGGTACAAGGCGATGATTCAATCATCTTGCTTATGTCCCATTATACTACAATCAAGGACTCATTTTTACCATTCTTCTCACTTTATGCTCAACAGTATTTTGGCTCTATAGTCAATACTAAGAAGAGCGAGGTGCTTCCTTCATTAGAAGGAGCTGAGGTACTAAAGTACCGCAACTTCGGCACTATGCCCAAGCGAGACGAACTACAACTATTAGCTATGTTAAGACATCCTGAGAGGACGTCTTCACTACCGTCCCTAATGGCACGAGCCATAGGAGTCGCCTACGCTAATTGTGGTAATCATACCCGTGTATACCAAATTTGCGAGGATATCCATAACTTCCTGGCTAAAGGCGGGTTTACCCCCGACCCTTATGGTTTACCAGGTAGTATTAGATACAGACAGAACTACATTCCAAGTTACGTGCAAGTTGACATTAGTCACTTTCCGACGTACTTTGAAACTGTCCAACATCTCCAAGATCCACAGCGTATTATCTTAACCGATAAGCACTGGCCTAAGGATCACTTTATCGGAATCCCCGGAAAGTCTTAAGTTTAAGACGTTATTTAGGCGTCACCTTAAGTTAATAAAATAAAAGGAAACCCCAGCTGCATGCAAAAAAAAAAAAAAAAATACT

>MsAPV2 586aa RdRp

MRNTTVIGHKPSLARPLYGNPDAGSNPAYADTVDHALRRYLTPEEFNRVVNGYRRSPWNEDALNDDIAKLNSDEHTVIKDEHYYKAIEHTKKLFTPKEKLRPVHFTDLRHYPWQLASNIGAPFASSKEWQHYVNDKFLKGHTATEVRDLFKEAHGQSLEPEIIDRRMTKRNLYNEMFLINRTNIHLIKDGRKTNDHGHDLRYWNTAFARQHLVEDDEPDKVRLVFGAPSTLLMAELMFIWPIQVSLLNRGPESPMLWGYETITGGWSRLHTWASTALPRFESVVTLDWSRFDKDARHTVIRDIHAMIMRPMFTFEHGYHPTYHYPDTSDTNPARLENLWNWMTDAILTTPLILPDGSVLRFNHSGIYSGYFQTQILDSIYNCVMIFTILSRMGFDLDRVEIKVQGDDSIILLMSHYTTIKDSFLPFFSLYAQQYFGSIVNTKKSEVLPSLEGAEVLKYRNFGTMPKRDELQLLAMLRHPERTSSLPSLMARAIGVAYANCGNHTRVYQICEDIHNFLAKGGFTPDPYGLPGSIRYRQNYIPSYVQVDISHFPTYFETVQHLQDPQRIILTDKHWPKDHFIGIPGKS

>MsAPV2 CP dsRNA2 1764nt

GACTCCGAATTGAATTGCATTCTATCTCGGTCTTATAAACTTAAAGAATTTTAGTCACAGCAGATTTTATTTTCTACTCCCCTCCCCCACATTATCATGGACAAATCTAAGAAGTATCTATCTGATGCTAAAGAAATCATCAAGGCTAATGGTCTTGAAGAAAAATTTCTCGCCGCTGCTGGTCTTTCATCTATGGATGAACTCGACCCTAAGAACTTCGCTGAGTTTGAAGCTCCTGCTCCTCCTCAAGTTATGAAGCCCTCTATCGCCGCTGCTCCCATCACTCAAAAACCGCTTTCCAAGCAGGTTGCTGATGCCTCTACTGGCTTTAATCCAGAGAGCGCTTCTGATATGCTCAGACCATTTATTGGTCTAAAGATTAACTACAAAGCTAGGAAGACTGTATCTACCTATGCTCCGTCCTCTCTTATGATGGATTATATTCTCCATCAAGTTAACAATTTACTTGTTGATAATTTCTACTTTCGTCGGGCTTGCCCCGATTACCACCCTTACATCTTACGCCTTTACTATGGCGTTTTGTTTTGGATTCAATGCATGCGTGCAATGAATCATGCTAGGCTGCTTCATGCTGAAGCCTCCCAATGTCTGTCAACATTCTTGGATGCCTTCCCTGCTGACTCCCTTCCAATTTCAAGTCCTTTACTTGACTTGTTCAAAACTCTTTGCTGCTCCCAACCAGAGATTCCCACTTTTGGAGTTGTATCTCCTACTCTCCCATTCTATCCTGGCCCTGAAGCTAGGAGCGCTTTTATGCTGGCTGATGCTACCAGCCATGTTTTACCCAACATTCCTGGGATTTTCGCTTTGATCGAGAACCTCCGGACCTGCCTCAACCCTGGCGATGGGCAACAGCCCAATCTGCCTGCTAAGGGTAAACATATCCCCGTTACCAGTACCGCCGCTGCGGCAACTGTTTTCGGTCACCACCAATTCCCAATTTTGGCCAACAGGTCAAATGCTGAAAAATGGAGCCTTTGCTCCTCTGGACTACAATATCCTTGTGAAGCCGATCGTCGACTTCACGAAAATTTTTCTGAAAGGCTCGATAACTTCGAGTTTCCTGTCACTGCCGAGGATGATGACCTCAGGTTTTATAGTGACTTTTTGCACATTACTAATAGTCTAGCTTGGTTTGCCCAAGTTAGAGATGTTGCTGCCAATGAAGCTGCTTTCTTCAATGGCTCTGGCACTTTGGCTGACTGTCCCCCTTTCGGGATTGCTGCCAATCAGATTACTGCGGGCTATTTAGCCCCAATTATTCAAGTTGCTGCTCCTACCCGGAGTGCCGACCCTGCCTCCCTCTTTCCATTCGCTATTAAGCTCGAAACCGCATCTCGAGCTCTTCCTGAATTAGCTGAAGCGATGGCTGCACTTGCGCAGACCAACGTGAATATCTACCCGACCCATCCCTACTACAACCACATTGGTGCTGAGTCGCGCCGTGGACCGTTCTGGAACCTTGCTCCCGTTGAATCCTCCATCACCGATCACTCGAGCTATCTCTCACTCCGAGAGGTTGTGCGCGGTTTAATGAAGCCCAAAGGCTAATTAAACTTGCCGGTTTTCATGTCTTCTTGTTTGAAGTTCTGATTTTCCGTTTTTCCTTTTTTGTTGAACCATTAATTGAAAACAAAAAAAAATATTTCCTTTGTTTAATTAAGGTTTTGAAGTAATTCTCTTCCCTTTTCGATTACCTCGTTTTTCGTTCTTAGTACTTTTATTTAAAAAAAAAAAAAAAAC

>CP 491aa

MDKSKKYLSDAKEIIKANGLEEKFLAAAGLSSMDELDPKNFAEFEAPAPPQVMKPSIAAAPITQKPLSKQVADASTGFNPESASDMLRPFIGLKINYKARKTVSTYAPSSLMMDYILHQVNNLLVDNFYFRRACPDYHPYILRLYYGVLFWIQCMRAMNHARLLHAEASQCLSTFLDAFPADSLPISSPLLDLFKTLCCSQPEIPTFGVVSPTLPFYPGPEARSAFMLADATSHVLPNIPGIFALIENLRTCLNPGDGQQPNLPAKGKHIPVTSTAAAATVFGHHQFPILANRSNAEKWSLCSSGLQYPCEADRRLHENFSERLDNFEFPVTAEDDDLRFYSDFLHITNSLAWFAQVRDVAANEAAFFNGSGTLADCPPFGIAANQITAGYLAPIIQVAAPTRSADPASLFPFAIKLETASRALPELAEAMAALAQTNVNIYPTHPYYNHIGAESRRGPFWNLAPVESSITDHSSYLSLREVVRGLMKPKG
